# Informatics Methods to Enhance Bacterial Peptide Detection

**DOI:** 10.64898/2026.10.06.757124

**Authors:** Bo Ren, Kenneth Weke, David R Goodlett, Javier A Alfaro

## Abstract

Detecting low-abundance bacterial peptides in samples dominated by human proteins using liquid chromatography–tandem mass spectrometry (LC–MS/MS) is challenging. Bacterial peptides are typically far less abundant than host-derived peptides, and sequence similarity between bacterial and human peptides can complicate confident attribution to a bacterial origin. To address this problem, we developed and benchmarked an *in silico* spectral library protein identification workflow for detecting *Staphylococcus aureus* USA300 peptides against a background of A375 cells. Peptide and protein identifications obtained at a 1% peptide-level false discovery rate were compared between the *in silico* spectral library search and a conventional sequence database search across a defined bacterial dilution series containing 0, 1%, 2%, 10%, 25%, 50%, and 100% bacterial protein. A human sequence similarity filter was applied to identifications from both search strategies to reduce the likelihood of retaining human-derived or human-like peptide sequences. Both search strategies showed concentration-dependent bacterial detection and shared a practical detection limit of approximately 1–2% bacterial protein but exhibited distinct performance profiles. The *in silico* spectral library search produced fewer bacterial-like background identifications in the human-only control, whereas MS²Rescore substantially increased spectral library identifications at higher bacterial concentrations, exceeding database search at 100% bacterial protein. The partially overlapping peptide sets identified by the two strategies indicate that they provide complementary coverage of the bacterial proteome.

## 1 Introduction

Proteomic detection of bacterial peptides by liquid chromatography–tandem mass spectrometry (LC–MS/MS) is challenging in human samples. Confident detection of bacterial proteins by proteomics can provide evidence of microbial presence that complements nucleic acid–based approaches by directly measuring microbial gene products^[1]^. Nucleic acid–based approaches, however, cannot readily distinguish viable bacteria from residual microbial DNA and are vulnerable to contamination in low-biomass samples^[2]^. In human samples suspected of containing bacteria, bacterial proteins may constitute only a minor fraction of the total measurable proteome. As a result, low-abundance bacterial protein can be masked by the much larger human-derived protein background^[3][4]^. In addition, sequence overlap between bacterial and human proteomes can make it difficult to confidently assign peptides to bacteria ^[3][5]^. These challenges highlight the need for workflows that evaluate both tandem mass spectral match quality and sequence distinguishability from the human proteome.

Conventional database search approaches remain the standard approach for peptide identification from LC–MS/MS proteomics data. Regardless of the database search software used, observed peptide tandem mass spectra are matched to candidate tandem mass spectra for peptides generated from protein sequence databases. The resulting peptide-spectrum matches (PSMs) are validated using a target–decoy false discovery rate (FDR) estimation^[6]^. This framework is powerful, but because of the aforementioned reasons it is not sufficient on its own to detect bacteria in the presence of a background of human peptides. The composition of the searched database determines the candidate peptide space, and changes in database breadth, redundancy, or taxonomic content can alter both sensitivity and the number of ambiguous or incorrect matches^[7][8][9][10]^. Moreover, an FDR estimated across all reported PSMs does not necessarily reflect the error rate within the small subset of putative bacterial identifications, particularly when bacterial-derived tandem mass spectra constitute only a small fraction of the total signal^[11]^. Consequently, a small number of high-scoring putative bacterial PSMs cannot be confidently interpreted as genuine bacterial identifications if comparable matches are also observed in human-only controls.

Spectral libraries are widely used in data-independent acquisition (DIA)^[12]^ workflows, which reduce stochastic precursor selection and improve the reproducibility of low-abundance peptide detection. In data-dependent acquisition (DDA), by contrast, fragmentation is biased toward the most intense precursor ions, such that low-abundance bacterial peptides may be sampled inconsistently or not at all. Although spectral library searching cannot recover precursors that were never selected for fragmentation, it may support the identification of acquired bacterial tandem mass spectra by incorporating expected fragment-ion patterns and relative intensities. We therefore evaluated whether an *in silico* spectral library could provide complementary bacterial peptide identifications from DDA data. The spectral library approach leverages the reproducibility of peptide fragmentation patterns by comparing an observed tandem mass spectrum with a reference tandem mass spectrum, including features such as the presence of fragment ions and their relative intensities. For bacterial detection, however, empirical spectral libraries generated from prior LC–MS/MS experiments may be unavailable or incomplete for the organism and biological condition of interest. Because these libraries contain only peptides observed during the experiments used for their construction, their coverage may be biased toward the proteins and peptides expressed and sampled under those specific conditions^[13][14][15]^. For example, the *E. coli* proteome detected directly in urine from patients with urinary tract infections (UTIs) has been shown to differ substantially from that of the corresponding isolates after just a single *in vitro* passage^[16]^. *In silico* spectral libraries address this omission by predicting peptide tandem mass spectra directly from a given genomic sequence, thereby enabling the generation of a custom bacterial library for the entire predicted proteome of a target organism without first acquiring a comprehensive empirical library, which might miss tandem mass spectra for select peptides ^[17][18][19]^. At the same time, predicted spectral libraries require careful evaluation because prediction error, expanded peptide search space, score calibration, and threshold selection can all contribute to false-positive matches. Therefore, *in silico* spectral library searches should be treated as a complementary approach that requires empirical benchmarking, rather than as an automatic replacement for database search or a traditional peptide spectral library.

Regardless of the approach used to assign peptide tandem mass spectra to a peptide sequence and by inference the protein from which it originated, degeneracy is problematic. For this reason, we prepared bacterial: human protein mixtures at 0:100, 1:99, 2:98, 10:90, 25:75, 50:50, and 100:0 (w/w) to benchmark bacterial peptide identification. These samples were used for benchmarking bacterial peptide detection because they provided both a known bacterial content and an empirical estimate of bacterial-like background.

Previously, such dilution-series experiments have been used to quantify the loss of bacterial peptides as bacterial biomass becomes a minor component of a mixed proteomic sample^[4]^. Although spectral library workflows are most commonly associated with DIA^[20]^, *in silico* spectral libraries have also been used to search DDA-MS data directly, including in metaproteomic benchmarks against conventional database search engines^[21]^. Studies show that dilution series and spectral libraries are useful, but they do not directly address how predicted *in silico* spectral library searches compare with conventional database searches for detecting low-abundance bacterial peptides in human samples suspected of containing bacteria. What remains less well defined is whether *in silico*-generated spectral libraries can yield bacterial identifications that are both above the bacterial-like background observed in human-only controls and sufficiently distinct from the human proteome. The specific contribution of this study is to combine, within a single benchmarking framework, a defined bacterial to human dilution series, human-only controls, human sequence similarity filtering, parallel database search, *in silico* spectral library search analyses, and empirical detection-limit estimation.

In this study, we used protein mixtures containing defined amounts of *Staphylococcus aureus* USA300 proteins and human A375 cell line proteins to create a dilution series for calibrating and benchmarking an *in silico*-generated spectral library search workflow aimed at improving the detection of low-level bacterial peptides in human samples. To do so, we generated an *in silico S. aureus* spectral library and used it to compare the *in silico* spectral library search approach with a conventional database search, both against mixtures ranging from 0% to 100% bacterial proteins. To reduce human-derived bacterial-like background, we applied a human sequence similarity filter and used human-only controls to empirically estimate false bacterial identifications. This design allowed us to evaluate sensitivity, specificity, method complementarity, and the practical detection limit of bacterial peptide identification. We then applied the calibrated workflow to test whether the detection limits defined by the dilution series could explain the bacterial identifications reported in a previous study of bacterially infected human cell lines.

## 2 Methods

### 2.1 Sample preparation

The A375 melanoma cell line was obtained from the American Type Culture Collection (ATCC). *Staphylococcus aureus* (USA300) was cultured in Tryptic Soy Broth (TSB) at 37°C with shaking at 200 rpm, overnight. Bacterial (USA300) and human cells were lysed separately using an ultrasonic probe to ensure efficient disruption of cellular membranes and release of intracellular proteins. Protein concentrations were determined using the Bradford assay. Based on the measured concentrations, bacterial and human protein lysates were mixed at defined ratios to generate a gradient series representing increasing bacterial content (0:100, 1:99, 2:98, 10:90, 25:75, 50:50, and 100:0, bacterial: human, w/w). Mixed protein samples were then subjected to standard bottom-up proteomic preparation. Proteins were precipitated using cold acetone, resuspended in urea buffer, reduced, alkylated, and digested with sequencing-grade trypsin overnight at 37 °C. Resulting peptides were purified using C18 solid-phase extraction prior to LC–MS/MS analysis.

### 2.2 Nano-LC–MS/MS analysis

Peptide samples were separated by online reversed-phase nano-liquid chromatography using an EASY-nLC 1000 system (Thermo Fisher Scientific). The system was equipped with an Acclaim PepMap 100 C18 trap column (100 µm × 2 cm, 5 µm, 100 Å) and an Acclaim PepMap 100 analytical column (75 µm × 150 mm, 3 µm, 100 Å), operated at a flow rate of 300 nL/min. The LC system was coupled online to an Orbitrap Fusion Tribrid mass spectrometer fitted with a Nanospray Flex NG ion source (Thermo Fisher Scientific). Mobile phase A consisted of 2% acetonitrile and 0.1% formic acid, and mobile phase B consisted of 90% acetonitrile with 0.1% formic acid. Peptides were separated using a 140-min gradient from 5% to 40% B, followed by a high-organic wash and re-equilibration. Mass spectrometric data were acquired in data-dependent acquisition (DDA) mode using Fusion Tune 3.5 software. Survey MS1 scans were acquired in the Orbitrap over an m/z range of 300–1500 at a resolution of 120,000 (at m/z 200), with internal calibration using the polysiloxane ion at m/z 445.12002. The instrument operated with a cycle time of 3 s, during which the most intense precursor ions were selected for tandem MS analysis. Precursors with charge states of 2–7 exceeding an intensity threshold of 5,000 were fragmented by higher-energy collisional dissociation (HCD) with stepped normalized collision energies of 28%, 30%, and 32%. Fragment ions were detected in the ion trap using a quadrupole isolation window of 1.6 Da. Dynamic exclusion was enabled to avoid repeated fragmentation of the same precursor ions.

### 2.3 Bacterial *in silico* spectral library generation

The reference proteome of *Staphylococcus aureus* strain USA300 was downloaded from UniProt (Taxon ID: 367830, 2026 January 5^th,^ 3,026 entries). This database was used to generate an *in silico* spectral library using MS2PIP (HCD2021 model)^[17]^. Protein sequences were subjected to *in silico* tryptic digestion, allowing up to 2 missed cleavages, and peptides of 8–25 amino acids were retained for downstream analysis. Spectra were predicted for tryptic peptides, assuming fixed carbamidomethylation of cysteine residues and variable oxidation of methionine residues. Decoy entries were also included during library generation to enable false discovery rate (FDR) estimation in downstream searches. The resulting target–decoy spectral library was subsequently formatted for spectral library searching. Detailed parameter settings for individual tools are provided in Supplementary Table A.

### 2.4 Database search

Raw LC-MS/MS data were analyzed using MSFragger^[22]^ within the FragPipe (v21.1) computational platform together with Philosopher (v5.1.0). Database search was performed against a *Staphylococcus aureus* USA300 protein database (Taxon ID: 367830). The USA300-only database was used intentionally to maintain the same bacterial proteome reference space across the complete 0–100% dilution series and to enable a direct comparison with the USA300 predicted spectral library. The objective of the search was not to profile the combined human and bacterial proteomes, but to evaluate the ability of each strategy to assign acquired spectra to candidate USA300 peptides. The human-only samples therefore served as empirical negative controls for bacterial-like assignments produced under this fixed search space. The default FragPipe workflow applies deep-learning-based rescoring of PSMs with MSBooster, followed by Percolator for PSM validation; thus, the database-search identifications were themselves refined by a machine-learning rescoring step. Detailed parameter settings for individual tools are provided in Supplementary Table B.

### 2.5 *in silico* Spectral library search

Raw LC–MS/MS data from samples containing 0–100% bacteria (USA300) were processed using the Trans-Proteomic Pipeline (TPP, v7.2.x)^[23]^. Vendor raw files were first converted to mzML format using the “Generate mzML” interface with default settings to ensure compatibility with downstream *in silico* spectral library searching. Spectral library searching was performed using the SpectraST Search module implemented in TPP. The processed USA300 spectral library was specified in the “Choose library file” field as the reference database for spectral matching. Candidate selection and scoring parameters were configured to detect homologous or identical hits up to rank 4, with carbamidomethylation (CAM) specified as the expected cysteine modification. All other parameters were left at default settings unless otherwise specified. Minimum F-value thresholds for reporting were set to 0.01 for top hits and 0.05 for lower-ranked hits, thereby retaining potential matches for subsequent filtering and threshold calibration.

To establish confident peptide identifications, peptide-level false discovery rate (FDR) control was applied. For each peptide sequence, only the highest-scoring spectrum match (by dot score) was retained as its representative identification, collapsing redundant spectrum matches to a single entry per peptide. These representative peptide identifications were then ranked in descending order of dot score, reflecting the spectral similarity between each peptide and its best-matching library entry. The FDR was estimated using a target–decoy approach, in which the cumulative ratio of decoy to target peptides was calculated along the ranked list. A q-value was assigned to each peptide as the minimum FDR observed at or below its dot score, and identifications were retained at a 1% FDR threshold. Peptides passing this threshold were considered confident identifications and carried forward for downstream analysis.

### 2.6 Peptide rescoring

As an alternative to the target–decoy peptide-level FDR filtering described in Section 2.5, peptide-spectrum matches (PSMs) from the *in silico* spectral library search were independently evaluated using a machine-learning-based rescoring strategy. These two approaches represent parallel confidence-filtering strategies applied to the same set of SpectraST search results: the traditional approach (Section 2.5) ranks identifications by the raw dot score, whereas the rescoring approach re-evaluates each PSM using additional predicted spectral features. This design enabled assessment of the spectral library search within a rescoring-based framework comparable to that used in the database search workflow (Supplementary Table C).

For rescoring, SpectraST search results were converted into the generic tab-separated format required by MS²Rescore (v3.2.1)^[24]^. Each PSM was represented by its peptidoform in ProForma notation (including precursor charge), the spectrum scan number, the raw file identifier, a target/decoy label inferred from the DECOY_ prefix, and the SpectraST dot score as the search-engine score; both target and decoy PSMs were retained without prior FDR filtering. Modifications were encoded using their Unimod names, with carbamidomethylation as a fixed modification on cysteine and oxidation as a variable modification on methionine.

Rescoring was performed with MS²Rescore using two complementary feature generators: MS²PIP (HCD2021 model) for predicted fragment-ion intensities and DeepLC^[25]^ for predicted retention times. Spectrum identifiers in the PSM file were matched to the corresponding mzML spectra via regular-expression patterns applied to the scan numbers. The resulting features, together with the original dot score, were used by the integrated Mokapot engine^[26]^ to train a semi-supervised model and to recalculate PSM-level scores and q-values. Peptide-level identifications were then derived as in Section 2.5, retaining peptides at a 1% FDR (Mokapot q ≤ 0.01).

Because the semi-supervised model requires a sufficient number of confident PSMs for training, rescoring could not be reliably executed for samples with very low bacterial content; for these samples, identifications were reported using the parallel target–decoy peptide-level FDR approach (Section 2.5). Peptides retained by either strategy were subsequently subjected to the same human sequence similarity ratio filter (Section 2.7).

### 2.7 Establishing Sequence Distinguishability: The Similarity Ratio

To quantify sequence similarity between candidate bacterial peptides and the human proteome, we defined a metric termed the similarity ratio. For each bacterial peptide, a self-alignment score was computed by aligning the peptide against itself, and a maximum human alignment score was obtained by aligning it against the most similar same-length (k-mer) peptide from the human reference proteome. The similarity ratio was then defined as the maximum human alignment score divided by the self-alignment score (best_human_score / self_score). All pairwise alignments were computed using the Biopython PairwiseAligner with the BLOSUM62 substitution matrix. To enforce ungapped alignments so that scores reflected residue substitutions only, gap-open and extension scores (including internal and end gaps) were set to a large negative value (−1×10⁶). The similarity ratio ranges from 0 to 1, where higher values indicate greater similarity to the human proteome and a value of 1 denotes a peptide identical to a human sequence. By scaling the human alignment score to each peptide’s own self-alignment score, the metric is normalized for peptide length and composition, enabling direct comparison across peptides of varying lengths.

A similarity ratio threshold of 0.75 was selected using the human-only and 100% USA300 calibration datasets following dot score–based filtering at 1% FDR (Supplementary Table D). At this threshold, all seven unique peptide identifications in the human-only control were excluded, whereas 1,993 of 2,386 unique peptides from the 100% USA300 samples were retained. Examples of excluded peptides are provided in Supplementary Table E. Peptides with a similarity ratio ≥ 0.75 were therefore classified as human-like and removed from downstream analysis (Figure 1). The same similarity ratio filter was applied identically to the database search results to ensure consistent removal of human-like sequences across both identification approaches.

**Figure 1.**
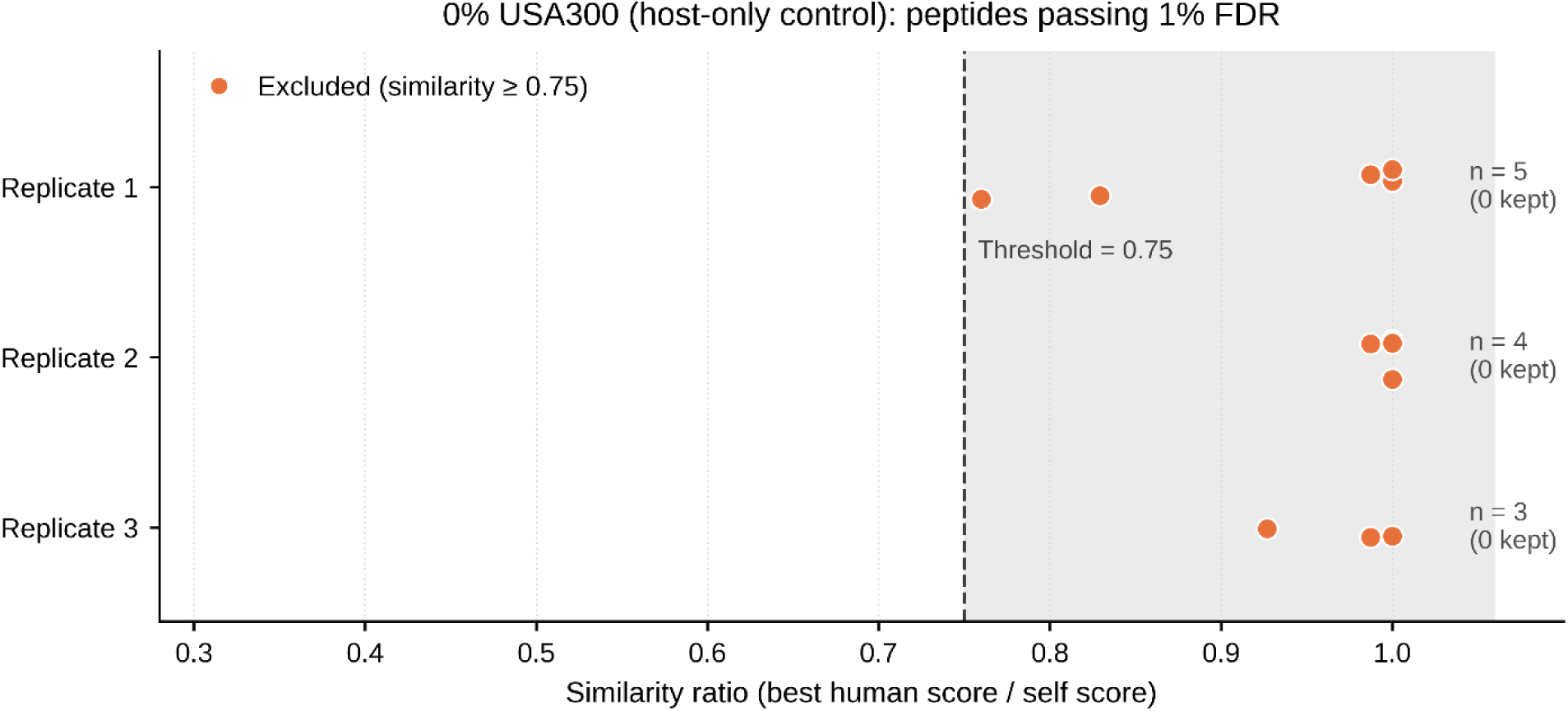
Similarity ratio distribution of peptides identified in the human-only (0% USA300) control across three replicates. Each point represents a unique peptide that meets the 1% FDR threshold in the *in silico* spectral library search, positioned according to its similarity ratio (the maximum human alignment score divided by its self-alignment score). Peptides at or above this threshold (orange, shaded region) were classified as human-like and excluded from downstream analysis. In this human-only control, all the identified peptides exhibited a similarity ratio ≥ 0.75, consistent with these identifications representing human-derived background rather than a bacterial signal.

An overview of the complete *in silico* spectral library search workflow, including the parallel target–decoy and rescoring-based confidence-filtering strategies and the subsequent human sequence similarity ratio filter, is provided in Figure 2.

**Figure 2.**
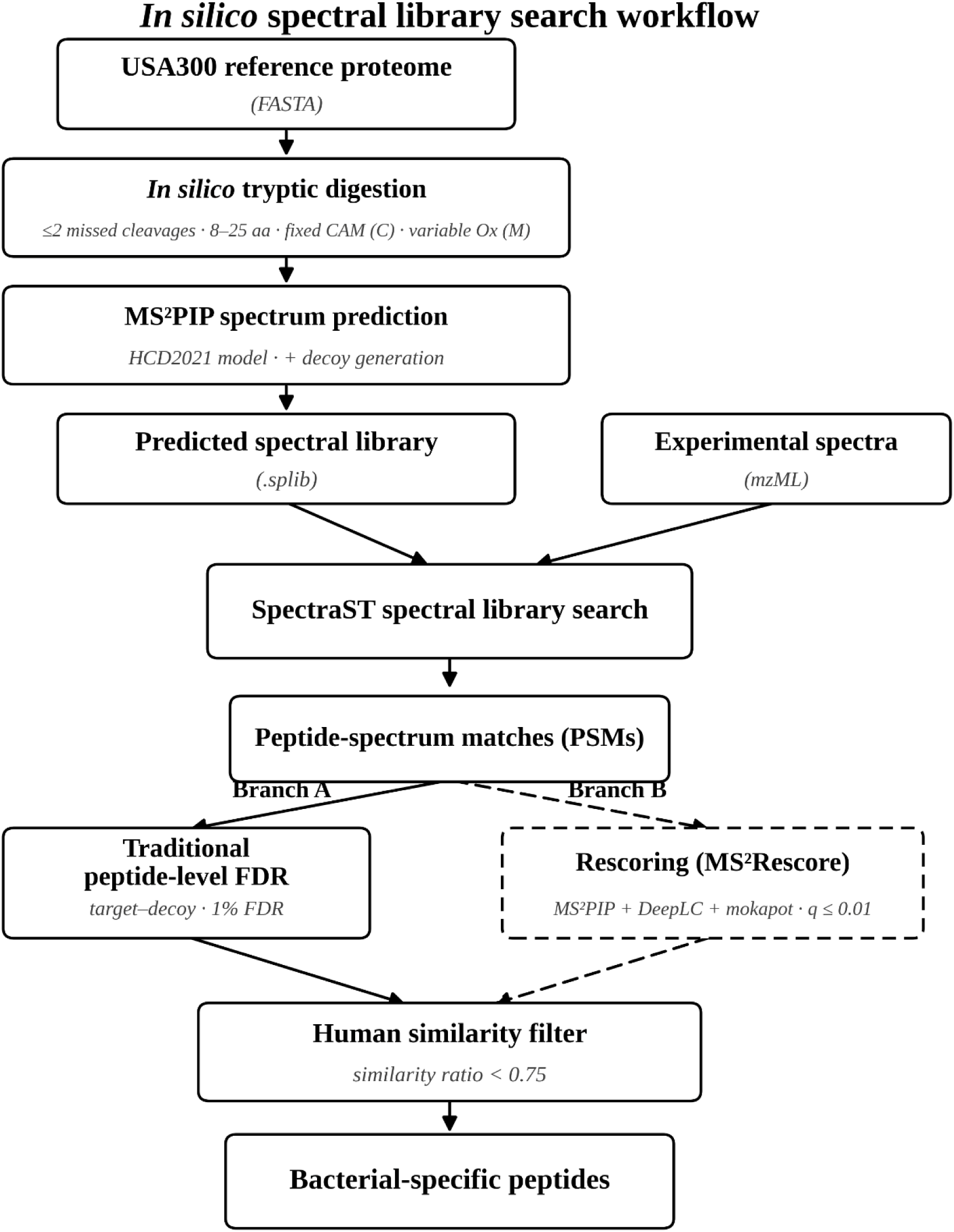
Overview of the *in silico* spectral library search workflow.

## 3 Results

### 3.1 Comparing *in silico* spectral library to database search across a USA300 dilution series

For both approaches, the bacterial candidate space was restricted to the USA300 proteome throughout the dilution series, allowing the human-only samples to provide an empirical measure of erroneous USA300 assignments under otherwise identical search conditions. Raw LC–MS/MS data from samples spanning 0%, 1%, 2%, 10%, 25%, 50%, and 100% bacteria were analyzed in parallel by *in silico* spectral library and database search, and confident identifications were retained at a 1% peptide-level FDR. Both approaches yielded concentration-dependent increases in the number of identified bacterial peptides and proteins (Table 1). At the 0% and 1% bacterial protein conditions, neither method produced reliable bacterial identifications after human sequence similarity filtering. Specifically, the *in silico* spectral library search identified no peptides in the human-only (0%) control and at most a few peptides at 1%, while the database search approach reported comparable background-level counts at 0% and 1% (17 and 16 unique peptides, respectively), indicating that these identifications largely reflected residual human background rather than genuine bacterial signal. Confident, reproducible detection emerged from the 2% sample onward and increased monotonically with bacterial content for both methods.

**Table 1.** Summary of unique peptides and proteins identified by *in silico* spectral library search and Database Search across a 0-100% bacteria (USA300) dilution series.

| % Bacteria (USA300) | Searching Method | Rep 1 (Unique Peptides) | Rep 2 (Unique Peptides) | Rep 3 (Unique Peptides) | Total Unique Peptides | Total Unique Proteins |
| --- | --- | --- | --- | --- | --- | --- |
| 0% (Human Only) | *In silico* spectral library Search | 0 | 0 | 0 | 0 | 0 |
|  | *In silico* spectral library Search+MS<sup>2</sup>Rescore | NA | NA | NA | NA | NA |
|  | Database search | 3 | 6 | 10 | 17 | 17 |
| 1% | *In silico* spectral library Search | 0 | 0 | 1 | 1 | 1 |
|  | *In silico* spectral library Search+MS<sup>2</sup>Rescore | NA | NA | NA | NA | NA |
|  | Database search | 3 | 7 | 9 | 16 | 14 |
| 2% | *In silico* spectral library Search | 4 | 4 | 4 | 9 | 8 |
|  | *In silico*<br>spectral library<br>Search+MS<sup>2</sup>Rescore | NA | NA | NA | NA | NA |
|  | Database search | 14 | 24 | 28 | 49 | 40 |
| 10% | *In silico*<br>spectral library Search | 42 | 49 | 57 | 69 | 40 |
|  | *In silico*<br>spectral library<br>Search+MS<sup>2</sup>Rescore | NA | NA | NA | NA | NA |
|  | Database search | 198 | 190 | 227 | 293 | 108 |
| 25% | *In silico*<br>spectral library Search | 276 | 227 | 232 | 344 | 125 |
|  | *In silico*<br>spectral library<br>Search+MS<sup>2</sup>Rescore | 461 | 398 | 447 | 606 | 186 |
|  | Database search | 585 | 545 | 527 | 776 | 200 |
| 50% | *In silico*<br>spectral library Search | 958 | 982 | 837 | 1175 | 373 |
|  | *In silico*<br>spectral library<br>Search+MS<sup>2</sup>Rescore | 1847 | 1758 | 1411 | 2142 | 546 |
|  | Database search | 1762 | 1736 | 1440 | 2179 | 476 |
| 100% | *In silico*<br>spectral library Search | 1553 | 1360 | 1692 | 1993 | 554 |
|  | *In silico*<br>spectral library<br>Search+MS<sup>2</sup>Rescore | 2457 | 2275 | 2534 | 3030 | 730 |
|  | Database search | 2099 | 2094 | 2324 | 2794 | 569 |

At the peptide level, database search identified more unique peptides than the unrescored *in silico* spectral library search at every bacterial concentration. The difference was greatest at 2% and 10% bacterial protein, where database search identified approximately 5.4- and 4.2-fold more peptides, respectively, and narrowed at higher bacterial concentrations to approximately 2.3-fold at 25%, 1.9-fold at 50%, and 1.4-fold at 100%. At the protein level, this difference narrowed with increasing bacterial content: at 50%, the two methods identified 476 (database search) and 373 (*in silico* spectral library search) proteins, and at 100%, they identified 569 and 554 proteins, respectively, indicating comparable proteome-level coverage at high bacterial content despite the lower number of peptides per protein identified by the *in silico* spectral library search. Notably, in the human-only control (0% bacteria), where any bacterial identification represents a false positive, the *in silico* spectral library search yielded no peptides after similarity filtering, compared with 17 for database search, indicating substantially fewer false-positive identifications for the *in silico* spectral library approach.

Because the database search workflow itself applies machine-learning rescoring, we re-evaluated the *in silico* spectral library peptide-spectrum matches with a comparable step (MS²Rescore), both to compare the two methods on equal footing and to test whether the lower peptide yield of the spectral library search reflected its simpler dot-score ranking rather than the spectral matching itself. For samples with sufficient bacterial signal to train the rescoring model (≥ 25%), rescoring substantially increased the number of confident bacterial peptides identified relative to the traditional target–decoy approach, by 58–81% across the 25%, 50%, and 100% conditions; e.g., 245→435 at 25%, 926→1,672 at 50%, and 1,535→2,422 unique peptides at 100%, mean of three replicates; Table 1. After rescoring, the number of unique peptides identified by the *in silico* spectral library search at 100% (3,030) exceeded that of the database search (2,794). In contrast, at low bacterial levels (≤10%) the number of confident identifications was insufficient to train the rescoring model, and these samples were reported using the traditional target–decoy approach. Rescoring, therefore, narrowed the peptide-level gap between the *in silico* spectral library search and the database search at high bacterial content, but did not extend detection below the ∼2% threshold imposed by the human background.

### 3.2 Overlap and complementarity of unique peptide identifications

To assess whether the two methods identified the same or distinct peptides, we compared their confident unique peptide identifications for each sample from 0% to 100% bacteria (Figure 3). At the lowest bacterial level (0–1%), the methods shared essentially no peptides, consistent with the lack of reliable bacterial detection at these levels. From 2% bacterial protein onward, the two methods showed increasing overlap in peptide identifications while also retaining sizable method-specific peptide sets. The number of shared peptides increased with bacterial content, reaching 1,188 at 100% bacterial protein (Figure 3G).

**Figure 3.**
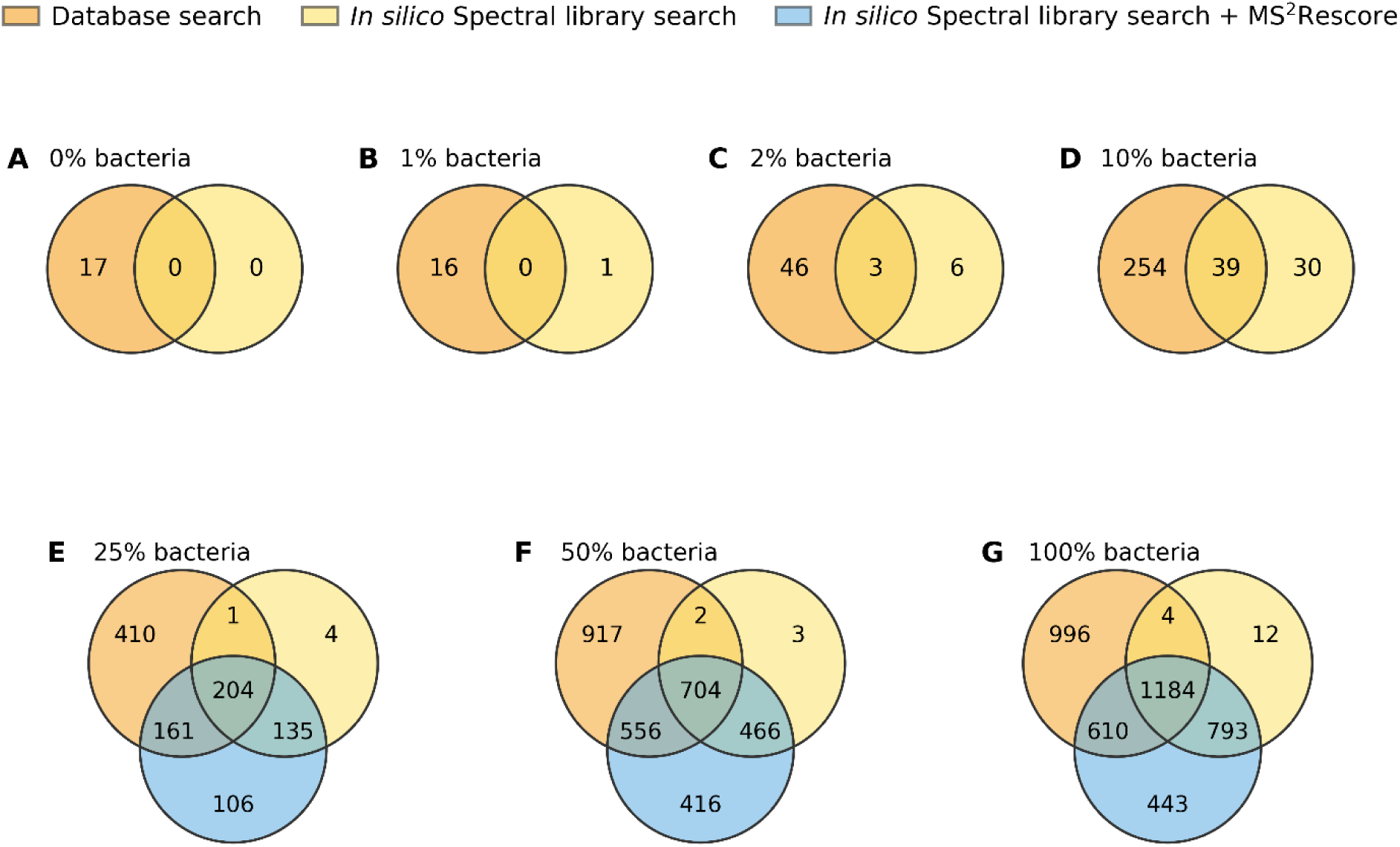
Complementarity among database search, *in silico* Spectral library search, and *in silico* Spectral library search with MS²Rescore across the 0% - 100% bacteria dilution series. Venn diagrams showing overlaps between unique bacterial peptides identified by database search and in *silico* spectral library search in samples containing (A-G) 0%-100% bacteria (USA300). Panels A–D (0%, 1%, 2%, and 10%) compare database search and *in silico* spectral library search; in panels E–G (25%, 50%, and 100%), where the bacterial signal was sufficient to support rescoring, the *in silico* spectral library search with MS²Rescore is included as a third set. Numbers indicate unique and shared identifications under the selected confidence threshold. At low abundance, the approaches detected largely distinct peptide subsets.

Consistent with its higher peptide-level sensitivity, the database search method contributed the larger set of exclusive peptides across all samples. Nevertheless, the *in silico* spectral library search identified a substantial number of peptides not detected by the database search method for all samples above the detection floor of ∼2%; for example, 139, 469, and 805 peptides exclusive to the *in silico* spectral library search at 25%, 50%, and 100% bacteria, respectively (Figure 3E–G). Since these identifications were retained under the same 1% peptide-level FDR applied to both methods, The method-exclusive peptide sets therefore support complementary coverage by the two search strategies.

For samples with sufficient bacterial signal to support rescoring (≥25%), we extended this comparison to a three-way analysis among database search, the *in silico* spectral library search, and the *in silico* spectral library search with MS²Rescore (Figure 3E–G). Rescoring retained nearly all peptides identified by the *in silico* spectral library search, while expanding both the shared set of peptides and the method-exclusive coverage. The set of peptides common to all three methods increased with increasing amounts of bacteria, reaching 204, 704, and 1,184 peptides at 25%, 50%, and 100%, respectively. Rescoring additionally identified a distinct set of peptides detected by neither database search nor the *in silico* spectral library search; i.e. 106, 416, and 443 rescoring-exclusive peptides at 25%, 50%, and 100%, respectively. Thus, MS²Rescore increased the overlap between the *in silico* spectral library search and the database search while continuing to contribute confident, method-exclusive identifications. Together, these results indicate that the *in silico* spectral library search and database search provide partially overlapping but substantially complementary coverage of the bacterial peptides, with the *in silico* spectral library search, particularly after MS²Rescore, contributing a distinct subset of confident identifications beyond those identified by database searching.

### 3.3 Performance in USA300-Infected Cell Models

To evaluate performance in a more authentic human–pathogen context, both search strategies were applied to A375 cells infected with *S. aureus* USA300 at multiplicities of infection (MOI) of 50:1 and 100:1 ^[27]^, and processed identically; i.e. 1% peptide-level FDR followed by the human sequence similarity filter. This analysis revealed that the *in silico* spectral library search identified essentially no bacterial-specific peptides, one peptide (one protein) at MOI 50:1 and none at MOI 100:1 (Table 2). As in the dilution series analysis, the few peptides initially passing the FDR threshold corresponded almost exclusively to highly abundant, conserved proteins (such as DnaK, MetK, ATP synthase subunits). Because these sequences closely resemble human homologs, they were removed by the human sequence similarity filter, leaving virtually no confident bacterial-specific identifications.

**Table 2.** Identification of USA300 peptides and proteins in infected A375 cell models.

| Infection Ratio<br>(USA300:A375) | Search Strategy | Peptide<br>number | Peptide<br>number<br>(unique) | Protein<br>number<br>(unique) | Common<br>Proteins |
| --- | --- | --- | --- | --- | --- |
| MOI 100:1 | Database Search | 3/6/7 | 10 | 4 | - |
|  | *in silico* Spectral<br>library search | 0/0/0 | 0 | 0 | - |
| MOI 50:1 | Database Search | 6/7/5 | 9 | 4 | 1 |
|  | *in silico* Spectral<br>library search | 0/0/1 | 1 | 1 | 1 |

The database search approach identified more peptides under the same workflow, 10 peptides (4 proteins) at MOI 100:1 and 9 peptides (4 proteins) at MOI 50:1 (Table 2), consistent with its higher peptide-level sensitivity. However, these counts were comparable to the false-identification background observed for database search in the human-only control (17 unique peptides at 0% USA300; Table 1), indicating that even database searching did not robustly exceed its own background at the bacterial load present in these samples. Increasing the MOI from 50:1 to 100:1 did not increase confident identifications for either method, and the overlap between the two methods was negligible (Table 2). In conclusion, these results show that in this intracellular infection model, neither approach achieved confident bacterial-specific detection clearly above background: the intracellular bacterial protein load was near the 1–2% detection limit defined by the dilution series, and the few bacterial proteins detectable at this load were predominantly conserved sequences that could not be confidently distinguished from the human proteome.

## 4 Discussion

### 4.1 Biological implications

To relate the dilution series analysis to what one might find on a biological scale, we estimated what a 1% bacterial protein fraction would represent at the cellular level. Using a HeLa-sized mammalian cell (∼150 pg total protein per cell) and an *E. coli*-like bacterial cell (∼100–234 fg protein per cell) as rough reference values^[28]^, a 1% bacterial protein fraction corresponds approximately to 6–15 bacterial cells per human cell. This calculation is only an approximation because protein mass varies across cell types, species, and physiological states. However, it indicates that even a seemingly low bacterial protein fraction can correspond to a substantial bacterial burden at the cellular level. In our dilution series data, clear separation from human-only controls did not appear until approximately 2% bacteria were present, indicating that bacterial signals can remain close to background even when bacterial content is already substantial on a per-cell basis.

### 4.2 Detection limit, specificity, and analytical trade-offs

A central difficulty in the study of human pathogens using proteomics is that pathogen-derived peptides are present at much lower abundance than human-derived peptides, so low-confidence background assignments can persist after conventional score filtering ^[10][29]^. A nominal 1% peptide-level FDR does not by itself demonstrate that a small number of USA300 assignments in a host-dominated sample represents genuine bacterial signal. Under the fixed USA300 candidate space used here, abundant human spectra may still produce high-scoring matches to bacterial peptide candidates, particularly when the true bacterial contribution is low. Human-only controls therefore provide an empirical estimate of erroneous USA300 assignments that complements target–decoy FDR control and helps determine whether putative bacterial signal exceeds the observed host-derived background ^[7][30]^.

Across the dilution series, database searching generally produced more peptide identifications, whereas the difference in protein-level coverage narrowed as bacterial content increased. This pattern suggests that the *in silico* spectral library search identified fewer peptides per protein but approached a similar breadth of bacterial proteome coverage at high bacterial abundance. The two measures therefore describe different analytical characteristics: peptide counts reflect identification depth, whereas protein-level coverage is often more directly relevant to biological interpretation in host–pathogen studies.

The two strategies also differed in their false-identification backgrounds. In the human-only control (0% bacteria), where any bacterial assignment is, by definition, a false positive, the *in silico* spectral library search yielded 0 bacterial peptides, compared with 17 for the database search. Under the fixed USA300 candidate space and calibrated human-similarity filtering conditions used here, the spectral library workflow produced a lower empirical background of erroneous USA300 assignments than the database-search workflow.

To test whether the peptide-level gap reflected a fundamental limitation of the *in silico* spectral library search or merely its simpler scoring, we re-evaluated the same SpectraST matches with machine-learning-based rescoring (MS²Rescore), which integrates predicted fragment-ion intensities and retention times within a semi-supervised model^[24]^. For samples with sufficient bacterial signal to train the model (≥25%), rescoring increased the number of confident bacterial peptides by 58–81% across the 25%, 50%, and 100% bacterial protein conditions (Table 1), so that at 100%, the rescored *in silico* spectral library search identified more unique peptides than database search (3,030 versus 2,794) and surpassed its protein-level coverage (730 versus 569 proteins). Importantly, rescoring preserved nearly all peptides identified by the dot-score workflow while substantially expanding the method-exclusive fraction (Figure 3E–G), indicating that the apparent sensitivity deficit of the *in silico* spectral library search was largely attributable to score calibration rather than to the underlying spectral matching. At low bacterial concentrations (≤10%), too few confident matches were available to train the semi-supervised model, so rescoring could not be applied and the gap persisted. Rescoring, therefore, improves sensitivity where a bacterial signal is already present but does not lower the fundamental detection limit imposed by the human background.

Consistent with this, both methods exhibited a similar lower limit of detection that rescoring did not overcome. At 1% bacteria, the number of bacterial peptides identified by database searching (16) was indistinguishable from its human-only background (17), and the *in silico* spectral library search identified at most a single peptide after human sequence similarity filtering, indicating that neither approach reliably detected bacterial signal at this level. Confident, concentration-dependent detection emerged only from 2% bacterial protein content onward, placing the practical detection boundary of the current workflow between 1% and 2% bacterial protein content. As noted in Section 4.1, this boundary already reflects a substantial per-cell bacterial burden, underscoring how severely human background constrains the detection of low-abundance bacteria regardless of search strategy or rescoring.

Even where the bacterial signal was abundant, the two search principles remained complementary rather than redundant. The *in silico* spectral library search identified a substantial set of confident peptides not detected by database search at every bacterial content above the detection limit (139, 469, and 805 method-exclusive peptides at 25%, 50%, and 100%, respectively; Figure 3E–G), and rescoring further identified peptides found by neither database search nor the dot-score workflow (106, 416, and 443 rescoring-exclusive peptides). Because all method-exclusive identifications were retained under the same 1% peptide-level FDR, they represent confident, complementary detections rather than low-confidence matches. Database search and the *in silico* spectral library search therefore occupy complementary points on a sensitivity–specificity trade-off: database search provides deep, well-optimized peptide-level coverage suited to high-confidence peptide identifications, whereas the *in silico* spectral library search offers comparable proteome coverage with a markedly lower false-identification background and a distinct subset of confident peptides. Two caveats temper this interpretation.

First, at low bacterial abundance, identifications from either method remain few and should be corroborated by reproducibility across replicates and, where possible, orthogonal validation such as targeted mass spectrometry. Second, the human sequence similarity filter, while effective at suppressing human-derived background, will also remove genuine bacterial peptides from highly conserved proteins, for example, chaperones of the HSP70/DnaK family, whose sequences closely resemble human counterparts; such conserved peptides therefore fall outside the detection scope of the filtered workflow.

### 4.3 Practical detection boundary

When applied to A375 cells infected with *S. aureus* USA300, the complete workflow identified few or no confident bacterial-specific peptides for either search strategy. This outcome is consistent with the detection limit established in the dilution series and reflects two compounding constraints. First, the intracellular bacterial protein load fell near the 1–2% lower limit of detection; increasing the MOI from 50:1 to 100:1 did not raise the number of confident identifications for either method, indicating that bacterial protein abundance, rather than acquisition depth, was limiting^[4]^ ^[20]^. Second, at this load the only bacterial proteins reaching detection were the most abundant ones, molecular chaperones, translation factors, and central metabolic enzymes, which are among the most evolutionarily conserved in the proteome and whose tryptic peptides are difficult or impossible to distinguish from human homologs at the sequence level^[31][32][33]^.

These constraints affected the two strategies differently but left both strategies unable to confidently detect bacteria above the background. For the *in silico* spectral library search, the small set of peptides initially passing the FDR threshold consisted almost entirely of conserved, abundant proteins, which were consequently removed by the human sequence similarity filter, leaving essentially no confident bacterial-specific identifications^[31][32]^. Database searching, owing to its higher peptide-level sensitivity, retained more peptides after filtering (9-10 peptides; 4 proteins), but this number was comparable to its false-identification background in the human-only control (17 unique peptides at 0% bacteria; Table 1), indicating that even database search did not robustly exceed its own background at this bacterial load. The apparent difference between the methods, therefore, reflects the higher baseline output, which has a correspondingly higher background, of database searching rather than a genuine detection advantage above noise.

We interpret these observations not as a deficiency in spectral library searching *per se*, but as a structural limitation of bottom-up proteomics for identifying trace levels of intracellular bacteria in a human sample. At this level, the detectable bacterial proteome is dominated by conserved sequences that cannot be confidently attributed to the pathogen rather than the human^[31][32][33]^. Because this constraint arises from bacterial abundance and sequence conservation rather than from the scoring or filtering algorithm, it is expected to apply broadly across search strategies. Overcoming it will likely require physical or biochemical enrichment of bacterial material prior to analysis ^[20][34]^, targeted assays directed at pre-selected, human-distinguishable bacterial peptides ^[4]^, or acquisition strategies with greater sensitivity for low-abundance precursors ^[20]^, rather than refinements of the search or filtering strategy alone. More fundamentally, where direct detection of the pathogen is limited in this way, as we demonstrated previously, the host response to infection may provide a more accessible indication of presence because bacterial “infection” induces reproducible changes in the host proteome^[27]^, offering sensitive, if indirect, evidence of infection even when bacterial peptides fall below the detection limit. Direct, pathogen-focused, and host-focused proteomics are therefore complementary, and the more informative choice depends on whether the goal is to identify and quantify the pathogen itself or to detect and characterize the infection.

## 5 Conclusion

We developed and benchmarked an *in silico* spectral library search workflow to detect low-abundance bacterial peptides in a background of a human cell line proteome, using a predicted *Staphylococcus aureus* USA300 spectral library with target–decoy false-discovery-rate control, and applying a human sequence similarity filter as a common downstream step for both the spectral library search and the database search. Across all samples from 0 to 100% bacteria analyzed at a 1% peptide-level FDR, both strategies detected bacterial signal in a concentration-dependent manner and shared a practical detection limit between 1% and 2% bacterial protein content. While database searching identified more unique peptides for each bacterial concentration, the two methods converged to comparable protein-level coverage at high bacterial content, and the unrescored *in silico* spectral library search exhibited a markedly lower empirical false-identification background in the human-only controls. The *in silico* spectral library search also identified a substantial, FDR-controlled set of confident peptides not detected by database search at any bacterial content above the detection limit, indicating genuinely complementary rather than redundant detections.

Applying machine-learning-based rescoring (MS²Rescore) to the spectral library matches substantially increased sensitivity wherever bacterial signal was sufficient to train the model (≥25% bacterial protein), raising peptide- and protein-level identifications above those of database search at high bacterial content (e.g., 3,030 versus 2,794 peptides and 730 versus 569 proteins at 100%). Because the model could not be reliably trained at bacterial concentrations of 10% or lower, the present data do not demonstrate that rescoring lowers the practical 1–2% detection boundary. Taken together, these results support the *in silico* spectral library search, particularly after rescoring, as a complementary strategy to database search that expands bacterial peptide coverage while sharing the same fundamental detection limit imposed by the human proteome background. The approach remains constrained by this low-abundance limit and by the exclusion of peptides from highly conserved, human-similar proteins, and its identifications, particularly at low bacterial abundance, warrant corroboration by reproducibility across replicates and orthogonal validation. Extending the workflow to authentic infection models, coupling it with bacterial enrichment to lower the detection limit, and characterizing the functional content of the complementary peptide set will further define the role of predicted spectral libraries in human–pathogen proteomics.

## Associated Data

The mass spectrometry proteomics data have been deposited to the ProteomeXchange Consortium via the PRIDE^[35]^ partner repository with the dataset identifier (PXD081839). The deposit also includes the analysis scripts used for peptide-level FDR estimation and for calculating the human sequence similarity ratio, as well as the search parameter and configuration files for all identification workflows.

## Acknowledgments

Work performed at the University of Victoria, Genome BC Proteomics Centre was supported by Genome British Columbia through funding to the BC Proteomics Centre for operations and technology development (374PRO).

## Declaration of Interest Statement

The authors declare that they have no known competing financial interests or personal relationships that could have influenced the work reported in this paper.

## Notes

### Competing Interest Statement

The authors have declared no competing interest.

## References

1. Do, K., Mehta, S., Wagner, R., Bhuming, D., Rajczewski, A. T., Skubitz, A. P. N., Johnson, J. E., Griffin, T. J., & Jagtap, P. D. (2024). A novel clinical metaproteomics workflow enables bioinformatic analysis of host-microbe dynamics in disease. mSphere, 9(6), e00793–23. 10.1128/msphere.00793-23

2. Eisenhofer, R., Minich, J. J., Marotz, C., Cooper, A., Knight, R., & Weyrich, L. S. (2019). Contamination in Low Microbial Biomass Microbiome Studies: Issues and Recommendations. Trends in Microbiology, 27(2), 105–117. 10.1016/j.tim.2018.11.003

3. Bakalarski, C. E., Elias, J. E., Villén, J., Haas, W., Gerber, S. A., Everley, P. A., & Gygi, S. P. (2008). The Impact of Peptide Abundance and Dynamic Range on Stable-Isotope-Based Quantitative Proteomic Analyses. Journal of Proteome Research, 7(11), 4756–4765. 10.1021/pr800333e

4. Timmins-Schiffman, E., Mikan, M. P., Ting, Y. S., Harvey, H. R., & Nunn, B. L. (2018). MS analysis of a dilution series of bacteria:phytoplankton to improve detection of low abundance bacterial peptides. Scientific Reports, 8(1), 9276. 10.1038/s41598-018-27650-4

5. Van Den Bossche, T., Kunath, B. J., Schallert, K., Schäpe, S. S., Abraham, P. E., Armengaud, J., Arntzen, M. Ø., Bassignani, A., Benndorf, D., Fuchs, S., Giannone, R. J., Griffin, T. J., Hagen, L. H., Halder, R., Henry, C., Hettich, R. L., Heyer, R., Jagtap, P., Jehmlich, N., … Muth, T. (2021). Critical Assessment of MetaProteome Investigation (CAMPI): A multi-laboratory comparison of established workflows. Nature Communications, 12(1), 7305. 10.1038/s41467-021-27542-8

6. Nesvizhskii, A. I. (2010). A survey of computational methods and error rate estimation procedures for peptide and protein identification in shotgun proteomics. Journal of Proteomics, 73(11), 2092–2123. 10.1016/j.jprot.2010.08.009

7. Blakeley-Ruiz, J. A., & Kleiner, M. (2022). Considerations for constructing a protein sequence database for metaproteomics. Computational and Structural Biotechnology Journal, 20, 937–952. 10.1016/j.csbj.2022.01.018

8. Tanca, A., Palomba, A., Fraumene, C., Pagnozzi, D., Manghina, V., Deligios, M., Muth, T., Rapp, E., Martens, L., Addis, M. F., & Uzzau, S. (2016). The impact of sequence database choice on metaproteomic results in gut microbiota studies. Microbiome, 4(1), 51. 10.1186/s40168-016-0196-8

9. Lee, E. M., Srinivasan, S., Purvine, S. O., Fiedler, T. L., Leiser, O. P., Proll, S. C., Minot, S. S., Deatherage Kaiser, B. L., & Fredricks, D. N. (2023). Optimizing metaproteomics database construction: Lessons from a study of the vaginal microbiome. *mSystems*, e00678–22. 10.1128/msystems.00678-22

10. Jagtap, P., Goslinga, J., Kooren, J. A., McGowan, T., Wroblewski, M. S., Seymour, S. L., & Griffin, T. J. (2013). A two-step database search method improves sensitivity in peptide sequence matches for metaproteomics and proteogenomics studies. PROTEOMICS, 13(8), 1352–1357. 10.1002/pmic.201200352

11. Wang, S., Feng, S., Pan, C., & Guo, X. (2022). FineFDR: Fine-grained Taxonomy-specific False Discovery Rates Control in Metaproteomics. 2022 IEEE International Conference on Bioinformatics and Biomedicine (BIBM), 287–292. 10.1109/BIBM55620.2022.9995401

12. Gillet, L. C., Navarro, P., Tate, S., Röst, H., Selevsek, N., Reiter, L., Bonner, R., & Aebersold, R. (2012). Targeted Data Extraction of the MS/MS Spectra Generated by Data-independent Acquisition: A New Concept for Consistent and Accurate Proteome Analysis. Molecular & Cellular Proteomics, 11(6), O111.016717. 10.1074/mcp.O111.016717

13. Lam, H., Deutsch, E. W., Eddes, J. S., Eng, J. K., King, N., Stein, S. E., & Aebersold, R. (2007). Development and validation of a spectral library searching method for peptide identification from MS/MS. PROTEOMICS, 7(5), 655–667. 10.1002/pmic.200600625

14. Lam, H., Deutsch, E. W., Eddes, J. S., Eng, J. K., Stein, S. E., & Aebersold, R. (2008). Building consensus spectral libraries for peptide identification in proteomics. Nature Methods, 5(10), 873–875. 10.1038/nmeth.1254

15. Deutsch, E. W., Perez-Riverol, Y., Chalkley, R. J., Wilhelm, M., Tate, S., Sachsenberg, T., Walzer, M., Käll, L., Delanghe, B., Böcker, S., Schymanski, E. L., Wilmes, P., Dorfer, V., Kuster, B., Volders, P.-J., Jehmlich, N., Vissers, J. P. C., Wolan, D. W., Wang, A. Y., … Röst, H. (2018). Expanding the Use of Spectral Libraries in Proteomics. Journal of Proteome Research, 17(12), 4051–4060. 10.1021/acs.jproteome.8b00485

16. Nartey, L. K., Weke, K., Yuen, V., Kibsey, P., Chen, M. X., & Goodlett, D. R. (2026). Proteomic Adaptations of *Escherichia coli* in Urinary Tract Infection Patients. Journal of Proteome Research, 25(3), 1714–1723. 10.1021/acs.jproteome.5c01165

17. Degroeve, S., Maddelein, D., & Martens, L. (2015). MS^2^ PIP prediction server: Compute and visualize MS^2^ peak intensity predictions for CID and HCD fragmentation. Nucleic Acids Research, 43(W1), W326–W330. 10.1093/nar/gkv542

18. Gessulat, S., Schmidt, T., Zolg, D. P., Samaras, P., Schnatbaum, K., Zerweck, J., Knaute, T., Rechenberger, J., Delanghe, B., Huhmer, A., Reimer, U., Ehrlich, H.-C., Aiche, S., Kuster, B., & Wilhelm, M. (2019). Prosit: Proteome-wide prediction of peptide tandem mass spectra by deep learning. Nature Methods, 16(6), 509–518. 10.1038/s41592-019-0426-7

19. Cox, J. (2023). Prediction of peptide mass spectral libraries with machine learning. Nature Biotechnology, 41(1), 33–43. 10.1038/s41587-022-01424-w

20. Willems, P., Fels, U., Staes, A., Gevaert, K., & Van Damme, P. (2021). Use of Hybrid Data-Dependent and -Independent Acquisition Spectral Libraries Empowers Dual-Proteome Profiling. Journal of Proteome Research, 20(2), 1165–1177. 10.1021/acs.jproteome.0c00350

21. Rajczewski, A. T., Mehta, S., Wagner, R., Gabriel, W., Johnson, J., Do, K., Vintila, S., Wilhelm, M., Kleiner, M., Searle, B. C., Griffin, T. J., & Jagtap, P. D. (2026). Comparative performance of Scribe and database search engines in metaproteomic profiling of a ground-truth microbiome dataset. Journal of Proteomics, 322, 105549. 10.1016/j.jprot.2025.105549

22. Kong, A. T., Leprevost, F. V., Avtonomov, D. M., Mellacheruvu, D., & Nesvizhskii, A. I. (2017). MSFragger: Ultrafast and comprehensive peptide identification in mass spectrometry–based proteomics. Nature Methods, 14(5), 513–520. 10.1038/nmeth.4256

23. Deutsch, E. W., Mendoza, L., Shteynberg, D., Slagel, J., Sun, Z., & Moritz, R. L. (2015). Trans-Proteomic Pipeline, a standardized data processing pipeline for large-scale reproducible proteomics informatics. PROTEOMICS – Clinical Applications, 9(7–8), 745–754. 10.1002/prca.201400164

24. Buur, L. M., Declercq, A., Strobl, M., Bouwmeester, R., Degroeve, S., Martens, L., Dorfer, V., & Gabriels, R. (2024). MS^2^ Rescore 3.0 Is a Modular, Flexible, and User-Friendly Platform to Boost Peptide Identifications, as Showcased with MS Amanda 3.0. Journal of Proteome Research, 23(8), 3200–3207. 10.1021/acs.jproteome.3c00785

25. Bouwmeester, R., Gabriels, R., Hulstaert, N., Martens, L., & Degroeve, S. (2021). DeepLC can predict retention times for peptides that carry as-yet unseen modifications. Nature Methods, 18(11), 1363–1369. 10.1038/s41592-021-01301-5

26. Fondrie, W. E., & Noble, W. S. (2021). mokapot: Fast and Flexible Semisupervised Learning for Peptide Detection. Journal of Proteome Research, 20(4), 1966–1971. 10.1021/acs.jproteome.0c01010

27. Ren, B., Weke, K., Hardie, D., Goncheva, M. I., Hupp, T., Alfaro, J. A., Pětrošová, H., & Goodlett, D. R. (2025). Proteomic Changes in Cancer Cell Lines as a Result of Bacterial Infection. *PROTEOMICS*, e70062. 10.1002/pmic.70062

28. Finka, A., & Goloubinoff, P. (2013). Proteomic data from human cell cultures refine mechanisms of chaperone-mediated protein homeostasis. Cell Stress and Chaperones, 18(5), 591–605. 10.1007/s12192-013-0413-3

29. Yang, Y., Hu, M., Yu, K., Zeng, X., & Liu, X. (2015). Mass spectrometry-based proteomic approaches to study pathogenic bacteria-host interactions. Protein & Cell, 6(4), 265–274. 10.1007/s13238-015-0136-6

30. Fels, U., Gevaert, K., & Van Damme, P. (2017). Proteogenomics in Aid of Host–Pathogen Interaction Studies: A Bacterial Perspective. Proteomes, 5(4), 26. 10.3390/proteomes5040026

31. Hao, C., Elias, J. E., Lee, P. K. H., & Lam, H. (2023). metaSpectraST: An unsupervised and database-independent analysis workflow for metaproteomic MS/MS data using spectrum clustering. Microbiome, 11(1), 176. 10.1186/s40168-023-01602-1

32. Holstein, T., Verschaffelt, P., Van Den Bossche, T., Van De Vyver, S., Martens, L., Mesuere, B., & Muth, T. (2026). The Peptonizer2000: Bringing Confidence to Metaproteomics. Journal of Proteome Research, 25(4), 1831–1845. 10.1021/acs.jproteome.5c00567

33. Nebauer, D. J., Pearson, L. A., & Neilan, B. A. (2024). Critical steps in an environmental metaproteomics workflow. Environmental Microbiology, 26(5), e16637. 10.1111/1462-2920.16637

34. Hoffmann, E., Machelart, A., Song, O.-R., & Brodin, P. (2018). Proteomics of Mycobacterium Infection: Moving towards a Better Understanding of Pathogen-Driven Immunomodulation. Frontiers in Immunology, 9, 86. 10.3389/fimmu.2018.00086

35. Perez-Riverol, Y., Bandla, C., Kundu, D. J., Kamatchinathan, S., Bai, J., Hewapathirana, S., John, N. S., Prakash, A., Walzer, M., Wang, S., & Vizcaíno, J. A. (2025). The PRIDE database at 20 years: 2025 update. Nucleic Acids Research, 53(D1), D543–D553. 10.1093/nar/gkae1011

