## Supplementary material for "Informatics Methods to Enhance Bacterial Peptide Detection": Supplementary_Materials.pdf

**Supplementary Table A.** $MS^2PIP$ *in silico* spectral library generation parameters

| Parameter | Value |
|---|---|
| fasta_file | USA300.fasta (UniProt Taxon ID 367830, 3,026 entries) |
| model | HCD2021 |
| cleavage_rule | trypsin |
| semi_specific | false (fully tryptic) |
| missed_cleavages | 2 |
| min_length | 8 |
| max_length | 25 |
| charges | 2, 3 |
| fixed modification | Carbamidomethyl (C) |
| variable modification | Oxidation (M) |
| max_variable_modifications | 3 |
| add_decoys | true |
| output format | MSP |

**Supplementary Table B.** MSFragger search parameters

### Mass Tolerance Settings

| Parameter | Value |
|---|---|
| precursor_mass_lower | -20 |
| precursor_mass_upper | 20 |
| precursor_true_tolerance | 20 |
| fragment_mass_tolerance | 20 |
| isotope_error | 0/1/2/3 |

### Enzyme and Digestion Settings

| Parameter | Value |
|---|---|
| search_enzyme_name_1 | stricttrypsin |
| search_enzyme_cut_1 | KR |
| search_enzyme_sense_1 | C |
| allowed_missed_cleavage_1 | 2 |
| digest_min_length | 7 |
| digest_max_length | 50 |
| digest_mass_range | 500.0 – 5000.0 |
| max_fragment_charge | 2 |

### Modification Settings

| Parameter | Value |
|---|---|
| variable_mod_01 | 15.9949 M 3 (Oxidation) |
| variable_mod_02 | 42.0106 [^ 1 (N-terminal Acetylation) |
| add_C_cysteine | 57.02146 (Carbamidomethylation, fixed) |
| max_variable_mods_per_peptide | 3 |
| max_variable_mods_combinations | 5000 |

### Output Settings

| Parameter | Value |
|---|---|
| output_format | pepXML_pin |
| output_max_expect | 50 |

### Ion Series

| Parameter | Value |
|---|---|
| fragment_ion_series | b, y |

**Supplementary Table C.** MS²Rescore configuration

| **Parameter** | **Value** |
|---|---|
| psm_file_type | tsv |
| feature generators | basic, ms2pip, deeplc |
| ms2pip model | HCD2021 |
| ms2pip ms2_tolerance | 0.05 |
| deeplc n_epochs | 20 |
| deeplc calibration_set_size | 0.15 |
| deeplc deeplc_retrain | false |
| rescoring engine | mokapot |
| psm_id_pattern | 0*(\d+) |
| spectrum_id_pattern | scan=(\d+) |
| lower_score_is_better | false |
| max_psm_rank_input | 10 |
| max_psm_rank_output | 1 |
| processes | 8 |

**Supplementary Table D.** Calibration of the human sequence similarity threshold

Peptide identifications passing the 1% peptide-level FDR threshold in the human-only control (A375, 0% bacteria) and in the 100% USA300 samples were scored against the human proteome, and the number retained was counted across a range of candidate similarity ratio thresholds. Counts are the union of three replicates. A threshold of 0.75 is the highest value at which no peptide from the human-only control is retained.

| Similarity ratio threshold | Peptides retained, human-only control (n = 7) | Peptides retained, 100% USA300 (n = 2386) | Retained (%), 100% USA300 |
|---|---|---|---|
| 0.60 | 0 | 1004 | 42.1 |
| 0.65 | 0 | 1364 | 57.2 |
| 0.70 | 0 | 1671 | 70.0 |
| 0.72 | 0 | 1810 | 75.9 |
| 0.75 | 0 | 1993 | 83.5 |
| 0.78 | 1 | 2116 | 88.7 |
| 0.80 | 1 | 2177 | 91.2 |
| 0.85 | 2 | 2312 | 96.9 |
| 0.90 | 2 | 2358 | 98.8 |
| 0.95 | 3 | 2376 | 99.6 |

**Supplementary Table E.** Peptides removed by the human sequence similarity filter (100% USA300)

Peptide identifications that passed the 1% peptide-level FDR threshold but were removed because their human sequence similarity ratio was at or above 0.75. Peptides with a ratio of at least 0.90 are listed here; the complete list of 565 removed peptides is available in the ProteomeXchange deposit. Peptides are pooled across the three replicates.

| **Peptide sequence** | **Similarity ratio** | **Protein** | **Removed in** |
|---|---|---|---|
| EAYPGDVFYLHSR | 1.000 | ATPA | both |
| ELIIGDRQTGK | 1.000 | ATPA | both |
| FTQAGSEVSALLGR | 1.000 | ATPB | both |
| FVIGGPQGDAGLTGRK | 1.000 | METK | MS2Rescore only |
| GFLLDGFPR | 1.000 | A0A0H2XH09 | MS2Rescore only |
| IGLFGGAGVGK | 1.000 | ATPB | both |
| NIGIMAHIDAGK | 1.000 | EFG | both |
| NIGIMAHIDAGKTTTTER | 1.000 | EFG | MS2Rescore only |
| TLLAQTLAK | 1.000 | CLPX | MS2Rescore only |
| VAELLLER | 1.000 | A0A0H2XGJ3 | both |
| IINEPTAAALAYGLDK | 0.987 | DNAK | both |
| VGIGPGSICTTR | 0.985 | IMDH | both |
| GVLLVGPPGTGK | 0.984 | A0A0H2XGW5 | both |
| VNIIDTPGHVDFTVEVER | 0.979 | EFG | MS2Rescore only |
| NAPCIIFIDEIDAVGR | 0.976 | A0A0H2XGW5 | both |
| IIIGGGMAYTFLK | 0.955 | PGK | MS2Rescore only |
| EADIDLLR | 0.947 | A0A0H2XFA9 | both |
| VALSINNK | 0.946 | A0A0H2XJZ9 | both |
| QTMLFSATMPK | 0.946 | CSHA | both |
| VFSFGPTFR | 0.939 | SYN | both |
| VSVLGHVQR | 0.932 | A0A0H2XGD7; PFKA | both |
| GGAIALGHPIGASGAR | 0.927 | THLA | both |
| LTEDIQGK | 0.925 | Y1685 | both |
| GLLVPVVK | 0.921 | A0A0H2XJG3 | both |
| LIVGGLEK | 0.921 | SYK | both |
| AIDALVPIGR | 0.917 | ATPA | both |
| SILINMIK | 0.917 | A0A0H2XHQ1 | both |
| TPPAPVLLK | 0.915 | RL11 | both |
| ILELSSSR | 0.912 | A0A0H2XFI1 | both |
| KGFLLDGFPR | 0.909 | A0A0H2XH09 | both |
| FIGNSYVK | 0.905 | A0A0H2XIJ3 | MS2Rescore only |
| NIDTGMGLER | 0.904 | SYA | MS2Rescore only |
| GAILVNAAR | 0.902 | A0A0H2XH62 | both |
| VAINGFGR | 0.902 | A0A0H2XHX6 | both |
| VKDLPGVR | 0.902 | RS12 | both |
| VILQDFTGVPAVVDLASLR | 0.900 | A0A0H2XKH5 | both |

**Supplementary Table F.** Peptide identifications in the infection model by *in silico* spectral library search

All peptide identifications passing the 1% peptide-level FDR threshold in A375 cells infected with *S. aureus* USA300, before and after the human sequence similarity filter. Replicates indicates the number of the three replicates in which the peptide was identified.

| MOI | Peptide sequence | Protein | Best dot score | Replicates | Similarity ratio | Outcome |
|---|---|---|---|---|---|---|
| 100:1 | IINEPTAAALAYGLDK | DNAK | 0.846 | 3 | 0.987 | Removed |
| 100:1 | FTQAGSEVSALLGR | ATPB | 0.760 | 3 | 1.000 | Removed |
| 100:1 | IGLFGGAGVGK | ATPB | 0.743 | 3 | 1.000 | Removed |
| 100:1 | FVIGGPQGDAGLTGR | METK | 0.700 | 2 | 1.000 | Removed |
| 100:1 | EAYPGDVFYLHSR | ATPA | 0.669 | 1 | 1.000 | Removed |
| 100:1 | YAIDETQR | A0A0H2XGZ8 | 0.666 | 1 | 0.780 | Removed |
| 100:1 | LVGIVTSR | A0A0H2XHN7 | 0.644 | 1 | 1.000 | Removed |
| 50:1 | IINEPTAAALAYGLDK | DNAK | 0.843 | 3 | 0.987 | Removed |
| 50:1 | IGLFGGAGVGK | ATPB | 0.739 | 3 | 1.000 | Removed |
| 50:1 | FVIGGPQGDAGLTGR | METK | 0.724 | 3 | 1.000 | Removed |
| 50:1 | FTQAGSEVSALLGR | ATPB | 0.763 | 2 | 1.000 | Removed |
| 50:1 | LLVEWNEK | RSMG | 0.687 | 2 | 0.886 | Removed |
| 50:1 | EAYPGDVFYLHSR | ATPA | 0.684 | 2 | 1.000 | Removed |
| 50:1 | TVIDYYHR | A0A0H2XH44 | 0.667 | 1 | 0.826 | Removed |
| 50:1 | TVGSGVVTEIIK | EFTU | 0.664 | 1 | 0.643 | Retained |
